# Impact of Human Serum Proteins on Susceptibility of *Acinetobacter baumannii* to Cefiderocol: role of iron transport

**DOI:** 10.1101/2021.08.18.456922

**Authors:** Casin Le, Camila Pimentel, Fernando Pasteran, Marisel R. Tuttobene, Tomas Subils, Jenny Escalante, Brent Nishimura, Susana Arriaga, Aimee Carranza, Alejandro J. Vila, Alejandra Corso, Luis A. Actis, Marcelo E. Tolmasky, Robert A. Bonomo, María Soledad Ramírez

## Abstract

Cefiderocol is a siderophore antibiotic that co-opts iron transporters to facilitate cell entry. We show that genes related to iron uptake systems and resistance to β-lactams in *Acinetobacter baumannii* have altered expression levels in the presence of human serum, human serum albumin, or human pleural fluid. Cefiderocol MICs are also raised in the presence of the mentioned fluids. Clinical response in *A. baumannii* infections may be related to the interplay of these human factors.

---

Carbapenem-resistant *Acinetobacter baumannii* (CRAB), one of the most feared pathogens in healthcare settings, driving the Centers for Disease Control and Prevention (CDC) to categorize it as an “urgent threat” (1–5). As a consequence of *A. baumannii*’s capability to develop multidrug resistance (MDR), treatment strategies become extremely limited, with only a few active antibiotics (6–8).

The increasing number of CRAB infections translates into alarmingly high morbidity and mortality (2, 9–11). Consequently, numerous efforts are focusing on finding novel treatment options (12–19). Cefiderocol (CFDC), formerly known as S-649266, approved by the FDA in November 2019 to treat nosocomial pneumonia and urinary tract infections, is a hybrid molecule that consists of a cephalosporin component that targets cell wall synthesis and a catechol siderophore moiety that allows cell penetration by active ferric-siderophore transporters (20–24). This novel synthetic compound uses a “Trojan horse” strategy to improve antibiotic penetration and reach a high concentration at the target site (21, 22). CFDC’s primary targets are cell wall synthetizing proteins associated with β-lactam activity (PBP1 and PBP3) (25). Early research showed promising results when treating Gram-negative carbapenem-resistant infections (7, 26, 27). Recently, a study described a number of unfavorable outcomes in patients with pulmonary or bloodstream *A. baumannii* infections compared to other available therapies (7). Our goal was to determine if there is a molecular basis for this observation.

*A. baumannii* senses components of human fluids and responds by modifying its transcriptional and phenotypic profiles (28–32). Human serum albumin (HSA) as well as human pleural fluid (HPF) modulate the expression of genes associated with iron-uptake systems, biofilm formation, antibiotic resistance, and DNA-acquisition among others (29, 33, 34). In previous studies we observed that CRAB AB5075 genes associated with iron uptake systems were down-regulated when exposed to HPF and 0.2% HSA (29), while genes associated with β-lactam resistance were up-regulated in the presence of physiological concentrations of HSA and human serum (HS) (33, 34). The modification in expression of iron-uptake systems and β-lactam resistance genes could be responsible for variations in *A. baumannii* susceptibility to CFDC, and concomitantly for the unfavorable outcomes of patients with pulmonary or bloodstream infections. In this work we describe the effect of HS, HSA, and HPF on representative CRAB strains.

CRAB AB0057 and AMA16 are clinical strains that belong to different clonal complexes and harbor OXA-23 and NDM-1, respectively (35, 36). Quantitative RT-PCR (qRT-PCR) assays were carried out using total RNA extracted from cells cultured in lysogeny broth (LB) or LB supplemented with 4% HPF or 3.5% HSA, or cultured in HS (Fig. 1A and B), as previously described (29).

**Figure 1.**
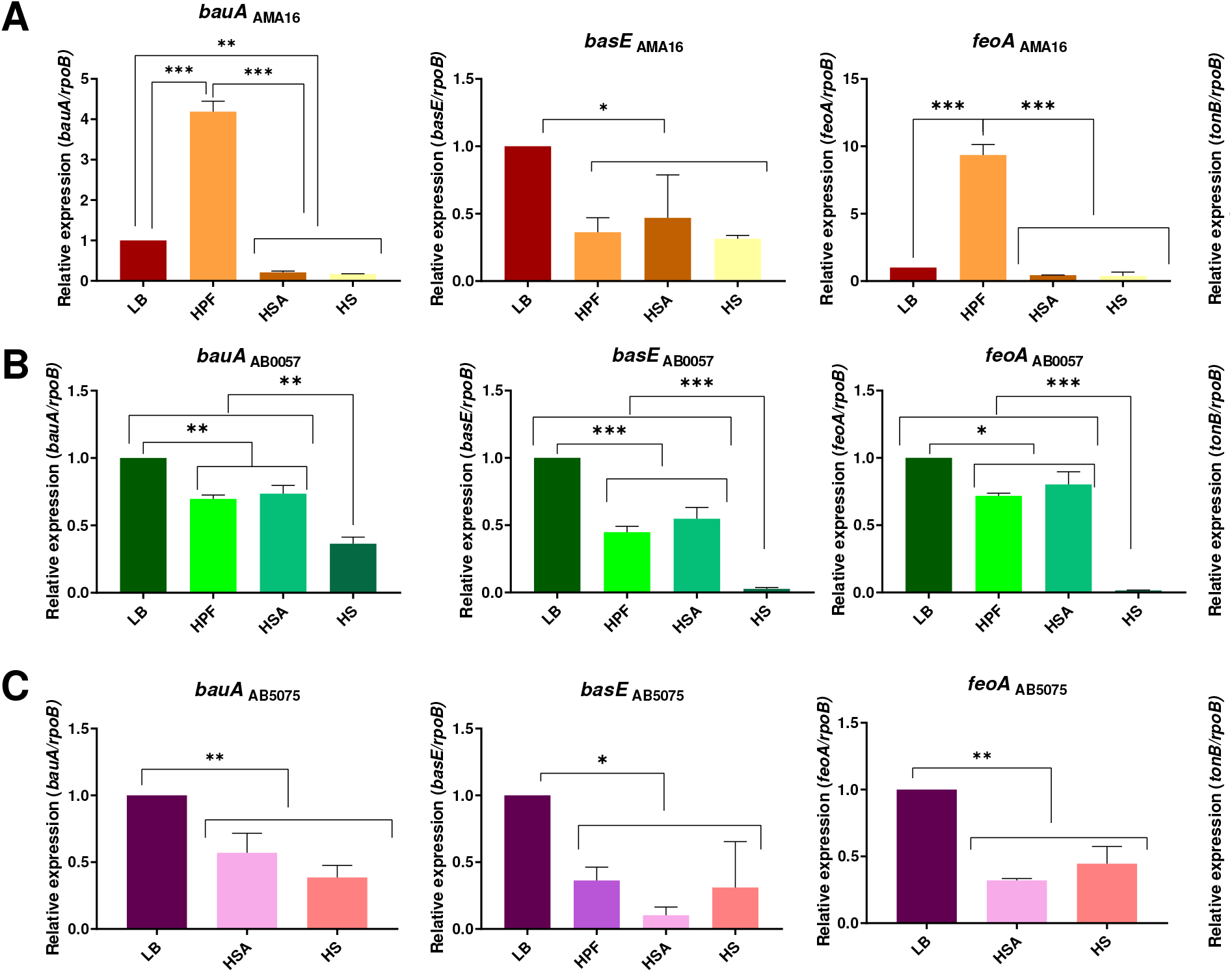
Genetic analysis of iron uptake genes of AMA16 (A), AB0057 (B) and AB5075 (C) *A. baumannii* strains. qRT-PCR of genes associated with iron uptake, *bauA*, *basE*, *feoA* and tonB expressed in LB or LB supplemented with HPF or HSA, or cultured in HS. Fold changes were calculated using double ΔCt analysis. At least three independent samples were used, and four technical replicates were performed from each sample. The LB was used as reference. Statistical significance (*p*< 0.05) was determined by ANOVA followed by Tukey’s multiple-comparison test, one asterisks: *p*< 0.05; two asterisks: *p*< 0.01 and three asterisks: *p*< 0.001.

The expression of iron uptake related genes *bauA*, *basE*, *feoA* and *tonB*, was reduced in both strains under most of the conditions evaluated (Fig. 1 A and B). In addition, when the hypervirulent *A. baumannii* AB5075 strain was exposed to 3.5% HSA or HS, the expression levels of most genes analyzed were down regulated. Pimentel *et al*. previously showed that levels of expression of AB5075 iron related genes were down-regulated in cells cultured in iron-rich media (29); this finding is in agreement with level of expression of *basE* reported in this work (Fig. 1C). The *fhuE_1*, *fhuE_2*, *pirA* and *piuA* genes, which are associated with iron-uptake (37), were also down-regulated in the presence of HPF (Fig. S1 A). Moreover, the transcriptional levels of these genes were further analyzed by qRT-PCR for the three strains (AMA16, AB0057, and AB5075) in the three tested conditions (HFP, HS, and HSA). In most of conditions evaluated, levels of expression were down-regulated (Fig. S1 B).

Genes associated with β-lactam resistance, such as *bla*_OXA-51-like_, *bla*_OXA-23_, *pbp1* and *pbp3* (33, 34), were up-regulated in *A. baumannii* AB5075, AMA16, and AB0057 when cultured in the presence of 3.5% HSA. To further examine if changes in genes associated with β-lactam resistance are affected by HS or HPF in the CRAB strains, qRT-PCR was used to assess gene expression of *bla*_OXA-51-like_, *bla*_OXA-23_, *bla*_NDM-1_, *pbp1*, *pbp3*, and *carO* in cultures of strains AMA16 and AB0057 containing HS or HPF. This transcriptional analysis revealed a down-regulation in expression levels of *bla*_NDM-1_ in *A. baumannii* AMA16 cultured in the presence of HS or HPF (Fig. 2 A). In this strain, the levels of expression of *ISAba125*, *bla*_PER-7_, *pbp1*, and *pbp3* were up-regulated in the presence of HPF. HS also induced up-regulation of ISA*ba125* (Fig. 2 A). In the case of the AB0057 strain, we observed up-regulation of *bla*_OXA-23_, *bla*_OXA-51_, *carO*, *bla*_ADC_, *pbp1* and *pbp3* in cultures containing HPF. In cultures containing HS the former four genes were up-regulated (Fig. 2 B). Assessment of the changes in β-lactam resistance-associated *A. baumannii* AB5075 genes in the presence of HPF showed down-regulation of the *carO* and *bla*_GES-14_ genes (Fig. 2 C). On the other hand, the expression levels of *bla*_OXA-51_, *pbp1*, *pbp3* and *bla*_ADC_ were increased (Fig. 2 C).

**Figure 2.**
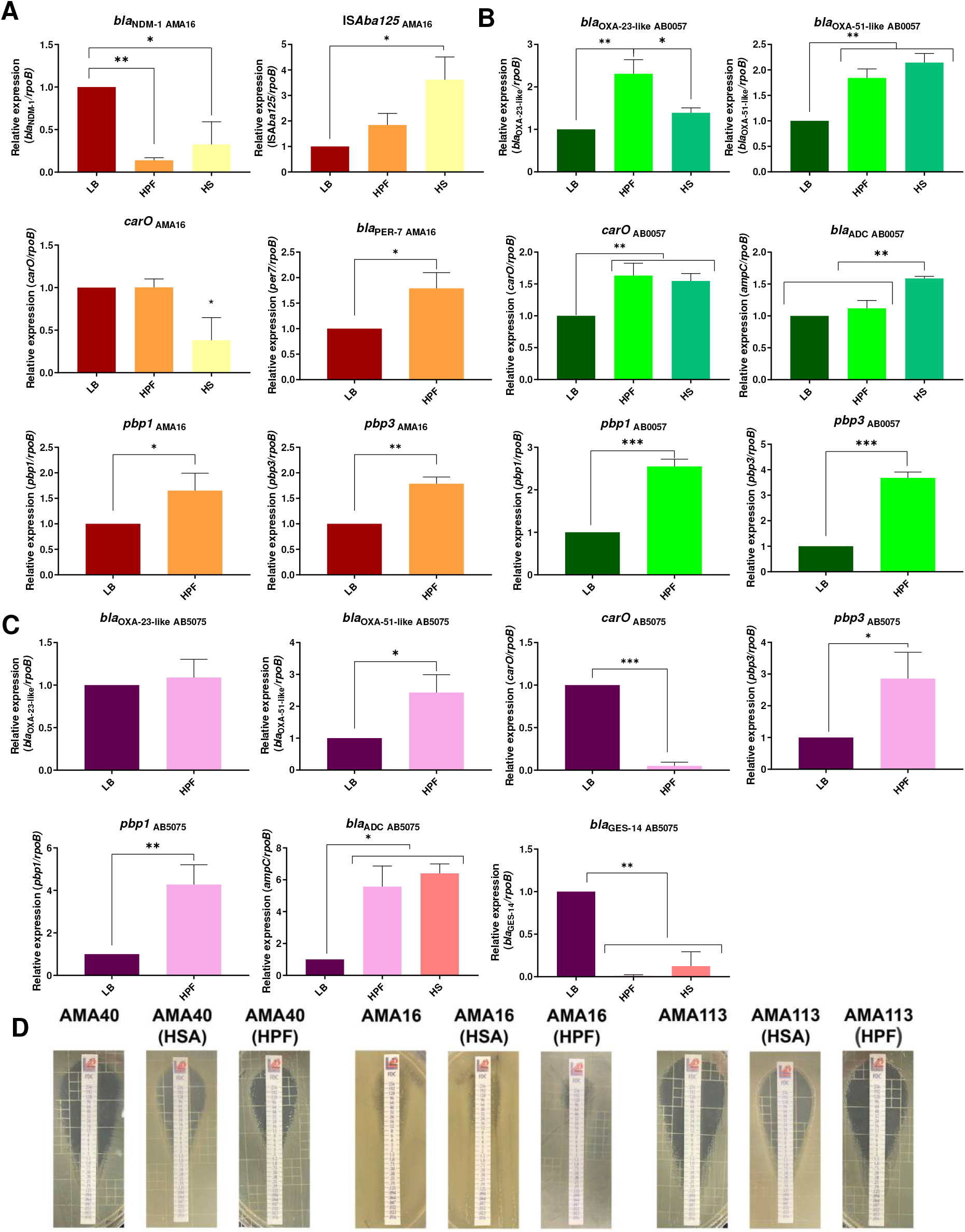
(A-C) Genetic analysis of β-lactamase + PBP genes of AMA16 (A), AB0057 (B) and AB5075 (C) *A. baumannii* strains. qRT-PCR of genes associated with β-lactams resistance expressed in LB, LB supplemented with HPF, or in HS. Fold changes were calculated using double ΔCt analysis. At least three independent samples were used. LB was used as the reference condition. Statistical significance (*p*< 0.05) was determined by ANOVA followed by Tukey’s multiple-comparison test, one asterisks: *p*< 0.05; two asterisks: *p*< 0.01, and three asterisks: *p*< 0.001. (D) Effect of HSA and HPF on the antimicrobial susceptibility of *A. baumannii* representatives strains AMA40 and AMA113 strains grew in LB broth, LB broth plus 3.5 % HSA, or HPF were used to performed cefiderocol (CFDC) susceptibility. Minimum inhibitory concentration (MIC) on cation adjusted Mueller Hinton agar was performed by MTS (Liofilchem S.r.l., Italy) following manufacter’s recommendations.

The impact of human fluids in CFDC activity was further studied using a panel of twenty CRAB strains including AB5075, AMA16, and AB0057 (a strain isolated in the Walter Reed Army Medical Center (38). Cells cultured in LB, and LB supplemented with HPF or 3.5 % HSA were used to determine the MICs of CFDC. Ten strains had baseline resistance to CFDC (Table S1). An increase in the MIC values was observed in 14 strains when the cells were growing in the presence of HSA or HPF (Table S1 and Fig. 2 D). We draw attention to the occurrence of heteroresistant colonies with the inhibition elipse in 10 of the tested strains (Fig. S2). Most of the strains that exhibited this phenomenon harbored *bla*_PER-7_ (35). Our results agree with a recent report indicating that PER-like β-lactamases contribute to a decreased susceptibility to CFDC in *A. baumannii* (39). In addition, they observed that the combination of CFDC with avibactam may inhibit the activity of PER-type ß-lactamases (39).

Overall, our results showed that *A. baumannii* responds to the interaction with human bodily fluids during the establishment of infection modulating the expression of iron-uptake genes and β-lactam associated genes. We hypothesize that the presence of iron binding proteins in the human fluids is sensed by *A. baumannii*, which in turn responds by down-regulating expression of iron uptake system genes impacting CDFC activity. The particular changes in gene expression of the aforementioned group of genes, which is not evaluated in traditional susceptibility tests, may contribute to unfavorable outcomes associated with iron-rich environments. These results also raise questions regarding the expression of other factors that may contribute to challenges in overcoming infections by *A. baumannii*.

The limitations inherent to performing MICs in the presence of bodily fluids (protein binding) is recognized. We are also aware that comparing these analysis does not take into account pharmacokinetics and pharmacodynamics (PK/PD) parameters that usure drug efficacy (40, 41). Laslty, we appreciate that these data must also be evaluated in the context of outcomes of animal models of infection (42–44). Nevertheless, the impact on iron transport mechanisms uncovered in these observations merits further analyses.

## Acknowledgements

We are grateful to Dr. Michael R. Jacobs for his critical reading of the manuscript.

## Author Contributions

C.P., C.L., F.P., M.R.T., T.S., R.A.B, and M.S.R. conceived the study and designed the experiments. C.P., C.L., F.P., M.R.T., T.S., J.E., B.N., S.A., A.C., A.J.V., L.A.A., M.E.T., R.A.B. and M.S.R. performed the experiments and genomics and bioinformatics analyses. C.P., C.L., F.P., M.R.T., T.S., A.J.V., L.A.A., R.A.B., M.E.T. and M.S.R. analyzed the data and interpreted the results. R.A.B., M.E.T. and M.S.R. contributed reagents/materials/analysis tools. F.P., M.R.T., T.S., L.A.A., R.A.B., M.E.T. and M.S.R. wrote and revised the manuscript. All authors read and approved the final manuscript.

## Supplementary Materials

**Table S1:** Minimal Inhibitory Concentrations of Cefiderocol (CFDC) for 22 Carbapenem-resistant *Acinetobacter baumanii* representative strains performed using CFDC MTS strips ((Liofilchem S.r.l., Italy) on Mueller Hinton Agar (cation adjusted). *A. baumannii* cells were cultured in LB or LB supplemented with 3.5 % HSA or HPF, respectively.

**Figure S1:** Differential expression of *fhuE_1*, *fhuE_2*, *pirA* and *piuA* genes associated with iron-uptake obtained for *A. baumannii* AB5075 cultured in the presence of HSA and HPF.

**Figure S2:** Intra-colonies in *A. baumannii* AMA40 representative of the tested strains in the presence of HPF.

